# Distinct patterns of skeletal muscle wasting and mitochondrial changes following hypoxia and inflammation: associations with physiological phenotype

**DOI:** 10.64898/2026.10.21.759645

**Authors:** Angelos Gavrielatos, Orane Lerouley, Noé Brocker, Cécile Cottet-Rousselle, Cindy Tellier, Clovis Chabert, Hervé Dubouchaud

**Affiliations:** Univ. Grenoble Alpes, Inserm U1055, Laboratory of Fundamental and Applied Bioenergetics (LBFA), Grenoble, France

**Keywords:** skeletal muscle, hypoxia, pulmonary inflammation, mitochondrial bioenergetics, protein turnover

## Abstract

Skeletal muscle demonstrates a high degree of plasticity that may enable mitochondrial and protein turnover adaptations to environmental stressors such as hypoxia and inflammation. However, the effects of these stressors on skeletal muscle mitochondrial function and proteostatic pathways remain incompletely understood and may depend on muscle phenotype. Given the potential heterogeneity in skeletal muscle responses to these stressors, this study investigated their independent and combined effects on mitochondrial function and protein turnover pathways in oxidative soleus and glycolytic plantaris and assessed the consistency of these responses by comparison with a previous cohort. Adult male Wistar rats were exposed for four weeks to normobaric hypoxia and/or lipopolysaccharide-induced pulmonary inflammation. Mitochondrial respiration, enzymatic activities, markers of mitochondrial function and protein turnover and spontaneous locomotor activity (SLA) were assessed. Pulmonary inflammation predominantly affected soleus, triggering atrophy alongside impaired CI-dependent mitochondrial function, without changes in mitochondrial density. In contrast, plantaris exhibited only atrophy with no mitochondrial alterations in response to inflammation. Hypoxia induced atrophy selectively in plantaris. Notably, the changes following exposure to inflammation and/or hypoxia occurred independently of SLA alterations while the combination of these two stressors did not produce additive effects for most parameters. Comparison with our previous cohort revealed divergent responses that may have been influenced by differences in several physiological characteristics between the two cohorts. These findings demonstrate that hypoxia and pulmonary inflammation induce muscle-specific adaptations and suggest that physiological heterogeneity may contribute to variability in skeletal muscle vulnerability to chronic stress.

**NEW & NOTEWORTHY:** This study demonstrates that chronic exposure to pulmonary inflammation and hypoxia exert distinct, muscle-specific and non-additive effects on skeletal muscle. Pulmonary inflammation predominantly alters mitochondrial function in oxidative muscle, whereas hypoxia preferentially induces atrophy in glycolytic muscle. Our findings further suggest that physiological heterogeneity may contribute to differences in skeletal muscle susceptibility to chronic stress.

## INTRODUCTION

Skeletal muscle is a highly plastic tissue capable of adaptations to a wide range of physiological and pathophysiological stimuli. These adaptations are largely governed by the ability of muscle to dynamically regulate energy substrate metabolism, protein turnover, and mitochondrial function, which together ensure the maintenance of cellular homeostasis under stress conditions (1,2). Among numerous stressors, hypoxia and inflammation represent two major factors with profound effects on skeletal muscle function and plasticity (3–5).

Chronic exposure to hypoxia (> 4500 m altitude) can lead to atrophy in both oxidative and glycolytic muscles independently of hypophagia (6,7) and has been associated with suppression of mammalian target of rapamycin complex 1 (mTORC1) signaling, partly through Regulated in DNA Damage and Development 1 (REDD1) upregulation alongside activation of ubiquitin–proteasome and calpain pathways (7–10). However, hypoxia can also exert muscle-specific effects as it has been shown to induce hypophagia-independent atrophy in highly glycolytic extensor digitorum longus (EDL), but not in oxidative soleus (11). Despite the well-described effects on muscle mass regulation, the impact of hypoxia on skeletal muscle mitochondrial function remains less clearly defined. We have recently shown muscle-specific alterations in ADP-stimulated oxygen consumption and reactive oxygen species (ROS) emission in response to chronic hypoxia (12) while earlier studies report either reductions (13) or no change in mitochondrial respiration (14,15). As hypoxia is frequently accompanied by reduced physical activity (16,17), which can independently modulate mitochondrial responses (18), failure to control for this confounder may contribute to these discrepancies. Collectively, these observations highlight the need for studies examining the effect of hypoxia, independently of activity levels, on mitochondrial function and protein turnover across muscles with distinct metabolic phenotypes.

Another well-established contributor to skeletal muscle dysfunction is inflammation that may originate not only within skeletal muscle but also in other organs (19). For example, inflammatory signaling originating from the lungs may result in skeletal muscle dysfunction through spill-over of cytokines and/or other mediators (20). This is supported by experimental data showing that pro-inflammatory cytokines such as tumor necrosis factor-α (TNF-α) lead to a switch towards glycolysis in C2C12 myotubes through the concomitant activation of hypoxia-inducible factor-1α (HIF-1α), suggesting a complex cross-talk between inflammatory and hypoxic signaling pathways (21). TNF-α may also disrupt the myogenic process in C2C12 myotubes via induction of caspases (22) and long non-coding RNAs (23) while exposure to chronic pulmonary inflammation through repeated lipopolysaccharide instillations inhibits the overload-induced hypertrophy (24) and decreases the mitochondrial calcium retention capacity in oxidative soleus (12). However, despite these observations, the literature remains scarce regarding the impact of chronic pulmonary inflammation on mitochondrial function and protein turnover pathways in skeletal muscle.

Given the marked differences in substrate utilization, mitochondrial content and redox regulation between oxidative and glycolytic muscles (25), it is reasonable to hypothesize that mitochondrial and metabolic responses to these two stressors may also be muscle-specific, complicating the interpretation of their respective effects on muscle alterations. In addition, these two factors may also coexist in pathophysiological conditions such as chronic respiratory diseases, including chronic obstructive pulmonary disease (26) and coronavirus disease 2019 (27), whereby muscle dysfunction is often observed (28). Nonetheless, the effect of their combination on skeletal muscle function has rarely been studied despite the possibility of synergistic or antagonistic interactions between their respective pathways.

The primary aim of this study was to explore the effect of chronic pulmonary inflammation and chronic hypoxia, separately and in combination, on multiple aspects of mitochondrial function and markers of protein turnover pathways in skeletal muscles with either oxidative (soleus) or primarily glycolytic (plantaris) phenotype. Given that hypoxia can alter activity levels, we also measured spontaneous locomotor activity to account for this potential confounding factor.

Importantly, inter-individual variability in the physiological response and adaptation to hypoxia may also exist (29), highlighting the possibility that skeletal muscle responses to hypoxia may vary across individuals and cohorts. Therefore, comparison of independent cohorts, under otherwise matched conditions, may provide insight into the consistency of skeletal muscle responses to hypoxia and potential sources of between-cohort variability. Therefore, comparison of independent cohorts, under otherwise matched conditions, may provide insight into the consistency of skeletal muscle responses to hypoxia and potential sources of between-cohort variability. We thus investigated as a secondary aim whether the skeletal muscle responses to hypoxia were consistent between the cohort of the present study and that one from our recently published study.

## MATERIALS AND METHODS

### Animals

The study complied with the European Union Directive (2010/63/EU) on laboratory animal use and care and was approved by the local Ethics Committee (Cometh 12) that is affiliated with the laboratory. It was further authorized by the French Ministry of Higher Education, Research and Innovation (APAFIS #40584-2023013114463378). Forty-eight Wistar male rats (20-24 weeks old; 484 ± 45 g) were housed in individual cages to monitor their food intake. As all animals in our previous cohort were male Wistar rats, we also restricted the present study to males to facilitate comparison between the two cohorts.

Animals were sacrificed via exsanguination from the abdominal aorta after anaesthesia with intraperitoneal injection of ketamine and xylazine (80 mg/kg and 50 mg/kg of body weight, respectively).

To delineate the effect of pulmonary inflammation and hypoxia, rats were assigned into one of the four following groups for 28 days until sacrifice: normoxia control (NC), chronic hypoxia (H), chronic pulmonary inflammation (I), H + I (HI). An overview of the experimental design is provided in Figure 1.

**Figure. 1.**
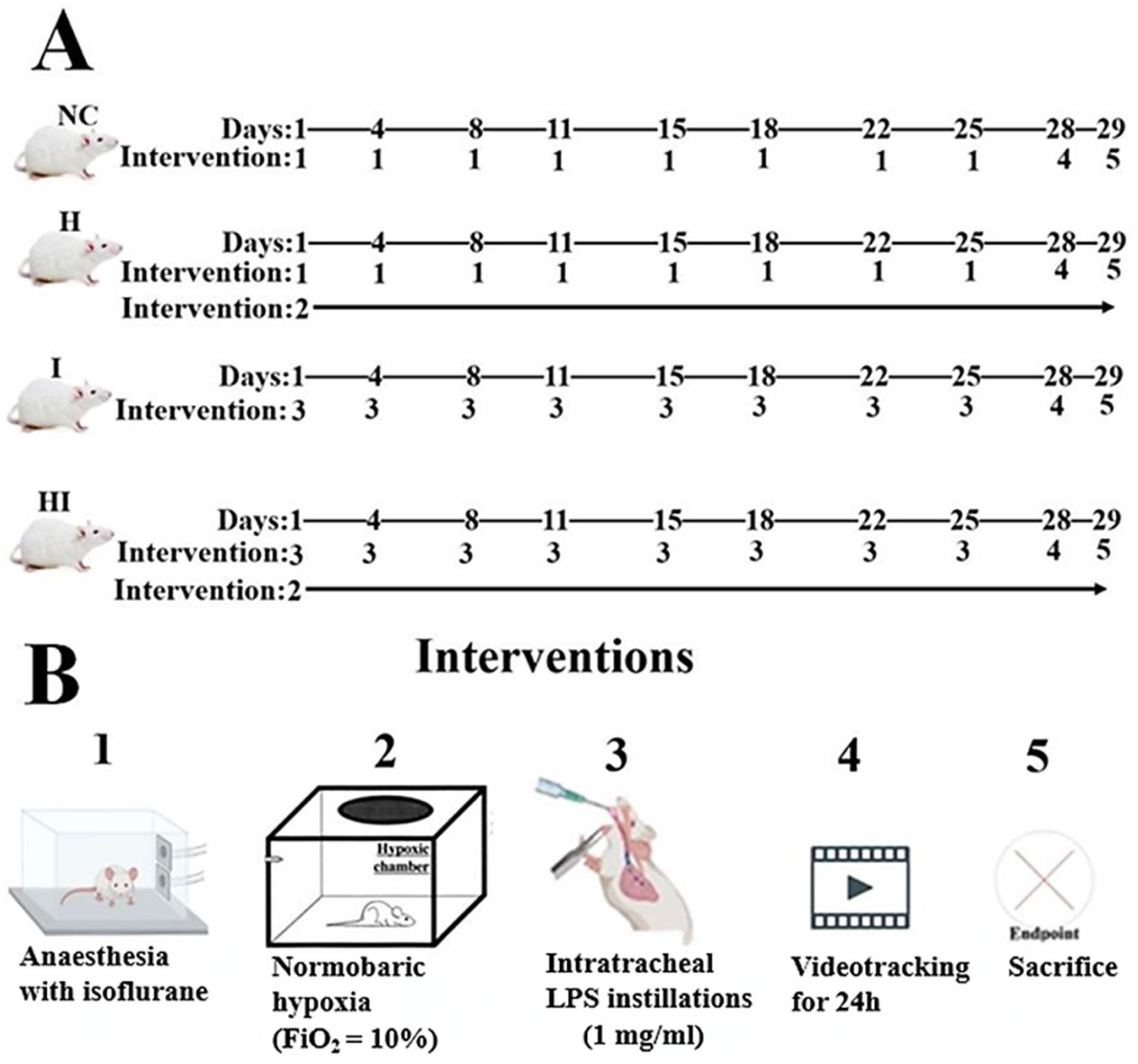
Experimental overview of the study. **A**: Timeline of all the interventions performed throughout the whole study period. **B**: Summary of the interventions performed.

### Pulmonary inflammation

To induce pulmonary inflammation to the rats from the I and HI groups, lipopolysaccharides (LPS; E. coli, serotype O55:B5; Sigma-Aldrich Chemical Co. ™, St Louis, Missouri, US in NaCl 0.9%) intratracheal instillations at 1 mg/kg were performed twice per week for four weeks. This chronic LPS administration protocol has been previously validated to induce pulmonary inflammation, resulting in structural alterations in rat lungs, including decreased alveolar area and increased alveolar wall thickness (30). The final instillation of LPS was performed at least 60 hours before sacrifice.

### Hypoxia

Animals from the H and HI groups were exposed to chronic continuous normobaric hypoxia for 4 weeks (Fraction of inspired O_2_: 10%). The hypoxic chambers were opened for 10-15 min every day to weigh the animals and their food pellets as well as to perform LPS intratracheal instillations in the rats from the HI group.

### Muscles sampling

Soleus and plantaris from the left hindlimb were dissected, weighed and snap frozen in liquid nitrogen before storage at -80°C until protein and gene expression analysis. Soleus and plantaris dissected from the right hindlimb were immediately transferred in an ice-cold isolation buffer (150 mM sucrose, 75 mM KCl, 1 mM KH_2_PO_4_, 5 mM MgCl_2_, 1 mM EGTA, and 50 mM Tris HCl, pH 7.4) to isolate the mitochondria.

### Heart dissection

Heart was excised and the right ventricle (RV) was separated from the left ventricle (LV) and interventricular septum (IVS). They were immediately weighted to determine the Fulton index as the ratio of RV/(LV+IVS) (31). This ratio is known to increase in response to chronic hypoxia due to pulmonary vasoconstriction and subsequent RV hypertrophy (32) and was thus used to validate the efficiency of our protocol or hypoxia exposure.

### Mitochondrial isolation

Soleus and plantaris were minced with scissors in ice-cold isolation buffer (150 mM saccharose, 75 mM KCl, 50 mM Tris base, 1 mM KH_2_PO_4_, 5 mM MgCl_2_, 1 mM EGTA, pH 7.4) supplemented with 0.2% fat-free bovine serum albumin (BSA). Minced muscles were incubated with subtilisin (0.24 mg/ml) for 1 min before new addition of isolation buffer and 0.2 % fat-free BSA. The muscle pieces were then homogenized using a motor-driven glass-Teflon pestle homogenizer. To remove cell debris and nuclei the suspension was centrifuged twice at 800 g for 10 min at 4°C. Mitochondria were isolated by spinning the resulting supernatant at 8000 g for 10 min at 4°C. The mitochondrial pellet was resuspended in respiration buffer containing (125 mM KCl, 5 mM Pi, 20 mM Tris HCl, 0.1 mM EGTA, and 0.1% fat-free BSA, pH 7.2) and kept on ice. Mitochondrial protein concentration was determined using the bicinchoninic acid assay (BCA, Pierce), with BSA as a standard.

### Spontaneous locomotor activity

Spontaneous locomotor activity (SLA) was recorded for 24 h, beginning 48 h before sacrifice using a digital camera (Switch Grip II, Samsung Electronics, Seoul, Republic of Korea) set to capture 1 picture/second. All video files were processed in MATLAB (v7.14, MathWorks r2012a, Natick, MA, US) as previously described (16). To evaluate SLA, we calculated the mean distance (cm/hour) covered throughout the entire videotracking period (24h) as well as separately during the dark (19:00 - 7:00) and the light phase (7:00-19:00).

### Oxygen consumption measurement

Rates of oxygen consumption were measured using a Clark type O_2_ electrode (Oxygraph; Hansatech Instruments, Pentney, UK). Mitochondria (0.2 mg/ml for soleus and 0.4 mg/ml for plantaris) were incubated at 30°C in respiration buffer and were stirred constantly with an electromagnetic stirrer. Mitochondria were energized using glutamate+malate (GM; 5 mM/2.5 mM) or succinate (S; 5 mM), either alone or in combination (GMS). Oxygen consumption was measured before (state 2) and after (state 3) the addition of 1mM adenosine diphosphate (ADP) and 1 μg/ml oligomycin (state 4). When succinate was used as a substrate alone, 4 µM rotenone were added to inhibit Complex I (EC 1.6.5.3). Mitochondrial respiration was normalized with citrate synthase (CS, EC:2.3.3.1) activity in the mitochondrial preparations. The quality of mitochondrial preparations was assessed using respiratory control ratios.

### Enzymatic activity assays

All the assays were performed at 30°C. CS activity was determined in whole muscle protein homogenates and in isolated mitochondria. Skeletal muscle homogenates (100 μg/ml) or isolated mitochondria (1 mg/ml) were incubated with 131.6 mM Tris-HCl, pH 8.1 supplemented with 0.09% Triton-X100, 150 mM DTNB and 100 µM Acetyl-CoA. Kinetics of release of sulphydryl and formation of mercaptide ion at 412 nm were recorded for 3 min with or without the presence of 100 µM oxaloacetate.

To determine CI activity, soleus homogenates (100 μg/ml) were incubated in a quartz cuvette at 30°C with 45.5 mM potassium phosphate buffer, pH 7.5 supplemented with 95 µM EDTA, 1 mg/ml BSA, 100 µM NADH and 100 µM decylubiquinone. NADH oxidation kinetics at 340 nm was recorded for 4 min with or without the presence of 10 µM rotenone.

The hydrolytic activity of ATP synthase (EC 7.1.2.2) was measured in soleus isolated mitochondria (130 μg/ml) incubated with 75 mM triethanolamine, pH 8.0, 5 mM MgCl_2_ and 20 µg/ml alamethicin. The reaction was initiated with addition of 4 mM ATP, and fixed volumes were collected at 0, 2, 4, and 6 min into a solution containing 0.38 M H_2_SO_4_, 5 µM ammonium molybdate, and 29 µM FeSO_4_. The inorganic phosphate (Pi) released by ATP hydrolysis was quantified spectrophotometrically at 750 nm with the use of a standard curve. The same experiment was performed after a 2-min pre-incubation with 5 µg/ml oligomycin to assess the oligomycin-insensitive ATP hydrolysis flux. Pi quantities at all four time points were plotted against time and the slope of the linear regression was used to calculate the Pi rate production. The hydrolytic activity of ATP synthase was normalized to citrate synthase activity in the isolated mitochondrial preparations.

To assess hydroxyacyl-Coenzyme A dehydrogenase (HAD, EC 1.1.1.35) activity, muscle homogenates (20 μg/ml for soleus, 200 μg/ml for plantaris) were incubated with 47.5 mM potassium buffer, pH 7.5 supplemented with 200 µM NADH, 60 µM EDTA and 40 mM imidazole. Following initial measurement of non-specific activity for 3 min, total HAD activity was quantified by monitoring the decrease in the NADH absorbance after addition of 50 µM acetoacetyl-CoA for 3 additional minutes.

Finally, lactate dehydrogenase (LDH, EC 1.1.1.27) activity was determined in muscle homogenates (soleus: 1 mg/ml; plantaris 0.5 mg/ml), incubated with 100 mM Tris buffer, pH 7.1 and 15 mM NADH. Following initial measurement of non-specific activity, total LDH activity was quantified by monitoring the decrease in the NADH absorbance at 340 nm after addition of pyruvate.

### Western blot analysis

Soleus and plantaris muscle portions (30-35 mg) were homogenized using a TissueLyser (MM301, Retsch Technology, Haan, Germany) in RIPA buffer (50 mM Tris HCl, 150 mM NaCl, 1.0% (v/v) IGEPAL® CA-630, 0.5% (w/v) sodium deoxycholate, 1 mM EDTA, 0.1% (w/v) SDS and 0.01% (w/v) sodium azide) supplemented with proteases (cOmplete tablets, Roche, Bâle, Switzerland) and phosphatases inhibitors (HaltTM Phosphatase Inhibitor Cocktail. ThermoFischer Scientific, Waltham, MA, USA). The homogenized samples were then centrifuged at 10000 x g at 4°C for 15 min and the supernatants were collected. Protein concentration was determined using the BCA assay. For gel electrophoresis, 20-30 μg of protein extracts, diluted with Laemmli buffer (0.25 M Tris-HCl pH 6.8; 8% (w/v) SDS; 40% (v/v) glycerol; 0.4 M DTT and 0.02% (w/v) bromophenol blue) were heated for 5 min at 100 °C (or 50 °C for oxidative phosphorylation content analysis) and then separated by SDS-PAGE on either 10-15 % custom-made gels or 4–20% Mini-PROTEAN® TGX™ precast gels (Bio-Rad Laboratories B.V.,Veenendaal, The Netherlands) for 60 min at 150-200 V. Analytical information regarding the targeted proteins, antibodies names, dilution and manufacturer information are reported in Supplemental Table S1. For OPA1 analysis, gels were run at 120 V for 3 hours to separate the long and short forms of OPA1. Proteins were transferred to polyvinylidene fluoride membranes (Bio-Rad) by electroblotting at 100 V for 45-60 min in transfer buffer (192 mM Glycine, 0.01% SDS, and 20% (v/v) ethanol. pH 8.3,). Following transfer, PVDF membranes were soaked in Amido Black (Sigma Aldrich, St-Louis, MI, USA) for at least 1 min and destained in solution containing 10% acetic acid and 25% isopropanol. Membranes were saturated with 5% non-fat dry milk in 50 mM Tris, 150 mM NaCl, pH 7.4, 0.1% Tween 20 (TBST) for 1 hour at RT. Membranes were then incubated overnight with primary antibodies (Supplemental table S1) at 4°C. Following incubation with appropriate secondary antibodies for 1 hour at room temperature, blot signals were revealed using enhanced chemiluminescence in an imaging system (Image Quant LAS 4000, GE Healthcare, Chicago, IL, US). Image J software was used to quantify band intensities in arbitrary units. For pAkt^ser473^/Akt analyses, membranes were harsh stripped by incubation in 10 % SDS, 0.5 M Tris HCl, pH 6.8, 0.8% ß-mercaptoethanol at 50°C for 30 minutes. Target protein quantity was corrected for total protein content using the Amido Black staining of the whole lane and normalised between gels using a gel-specific internal reference. For most blots, this reference consisted of a pool of samples. For a few blots that had no pooled sample, the mean signal of the two NC samples included on the same gel was used as the internal reference.

### RT-qPCR

Total RNA was extracted after homogenization of 45–60 mg of soleus and plantaris muscle portions using the TissueLyser (MM301) in the presence of 1 ml TRIzol™ reagent (Invitrogen), using a phenol-chloroform protocol including two rounds of phase separation with chloroform, precipitation with isopropanol (100%), and three ethanol washes. RNA concentration and purity were assessed using a UV-Vis spectrophotometer (NanoDrop™, ThermoFisher Scientific ™, Waltham, Massachusetts, USA). Reverse transcription was performed using 1 μg of RNA with Superscript III kit (Invitrogen, Carlsbad, CA, US) according to the guidelines of the manufacturer. qPCR reactions contained SYBR Green Master mix (Brilliant III Ultra-Fast SYBR Green, Agilent, Santa Clara, CA, US), forward and reverse primers at their respective concentrations (Supplemental table S2) and RNase-free water. Hypoxanthine phosphoribosyl transferase (HRPT) was used as a housekeeping gene. Quantification of the PCR product was performed either with the Livak method (E**^−ΔΔCt^;** (33)) or with the Pfaffl method (34) depending on whether HRPT and each target gene exhibited the same PCR efficiency. To verify the specificity of the PCR amplification, a melting curve analysis was performed following PCR.

### Fiber typology analysis

As skeletal muscle fiber typology is a major determinant of muscle metabolic phenotype, we sought to compare the soleus fiber composition between the two cohorts and provide further insight into potential sources of between-cohort variability. We thus determined soleus fiber typology in control animals of the present study to compare it with the corresponding control animals of our recently published study. Soleus fiber typology was determined via immunofluorescent staining of myosin heavy chain (MHC) I, MHC IIa and laminin. Specifically, following incubation for 1h at 37°C with primary antibodies targeting MHC I (BA-F8 IgG2b, 1:50), MHC IIa (SC-71 IgG1, 1:100) and laminin (D18 IgG2a, 1:50), slides were washed with PBS and immediately incubated for 1h at 37°C with species- and subclass-specific fluorescent secondary antibodies (IgG2b Alexa Fluor 488, 1:300; IgG1 Alexa Fluor 568, 1:300; IgG2a Alexa Fluor 594, 1:300). Primary and secondary antibodies were obtained from the Developmental Studies Hybridoma Bank (University of Iowa, IA, USA) and Invitrogen (Waltham, MA, USA), respectively. Slides were mounted with ProLong Gold Antifade (Invitrogen) and cured overnight at room temperature under light-protected conditions.

Image acquisition analysis was performed as described previously (12) using a Leica DMi8 wide-field epifluorescence microscope (Leica Microsystems, Wetzlar, Germany). Fiber type classification was conducted manually using Fiji (v1.54m) and the abundance of each fiber type was calculated as a percentage of the total number of fibers.

### Statistical analysis

Group size was determined using the statistical software Sigma-Plot 14, based on previous studies from our group using the same or similar experimental models and with parameters including minimum detectable difference in markers of mitochondrial function and protein turnover pathways, expected standard deviation of residues and desired power. For reasons of data quality or predefined criteria of statistical quality control (removal of outliers), the number of samples analyzed varied between measurements. All these exclusions were applied prior to statistical analyses and were not based on the experimental outcome. Normality of the data was determined using the Shapiro-Wilk test and histograms/boxplots. Where data did not follow a normal distribution, log10, square root or reciprocal transformations were conducted. To examine the effect of the two experimental conditions (normobaric hypoxia and pulmonary inflammation) a 2-way analysis of variance (ANOVA) was performed with condition 1 (normoxia or hypoxia) and condition 2 (with or without inflammation) as between-subjects factors. In case of significant interactions (condition 1 x condition 2), then Bonferroni-corrected 1-way ANOVA and subsequently t-tests were employed for further pairwise comparisons. To understand whether between-cohort variation in the responses to the two stressors may be linked to differences in physiological and metabolic characteristics, we compared several physiological and biological traits (i.e., initial body weight, fiber typology) between the two cohorts using independent-samples t-tests or Mann–Whitney U tests. Pearson or Spearman correlation analyses were performed to examine associations between physiological characteristics and relevant response variables, and linear regression analyses using ordinary least squares were conducted to quantify potential associations. Data were expressed as mean and standard deviation (SD). Values deviating ≥ 2 SD from the mean were excluded prior to statistical analyses. Statistical significance was determined as p < 0.05. To quantify effect sizes partial eta square (η^2^_p_) values were used, with a η^2^_p_ value ≥ 0.14 indicating a large effect.

## RESULTS

### Spontaneous locomotor activity

There was no difference in the distance covered per hour recorded two days before sacrifice, either during the night or day or throughout the entire 24-hour period (Figure 2).

**Figure 2.**
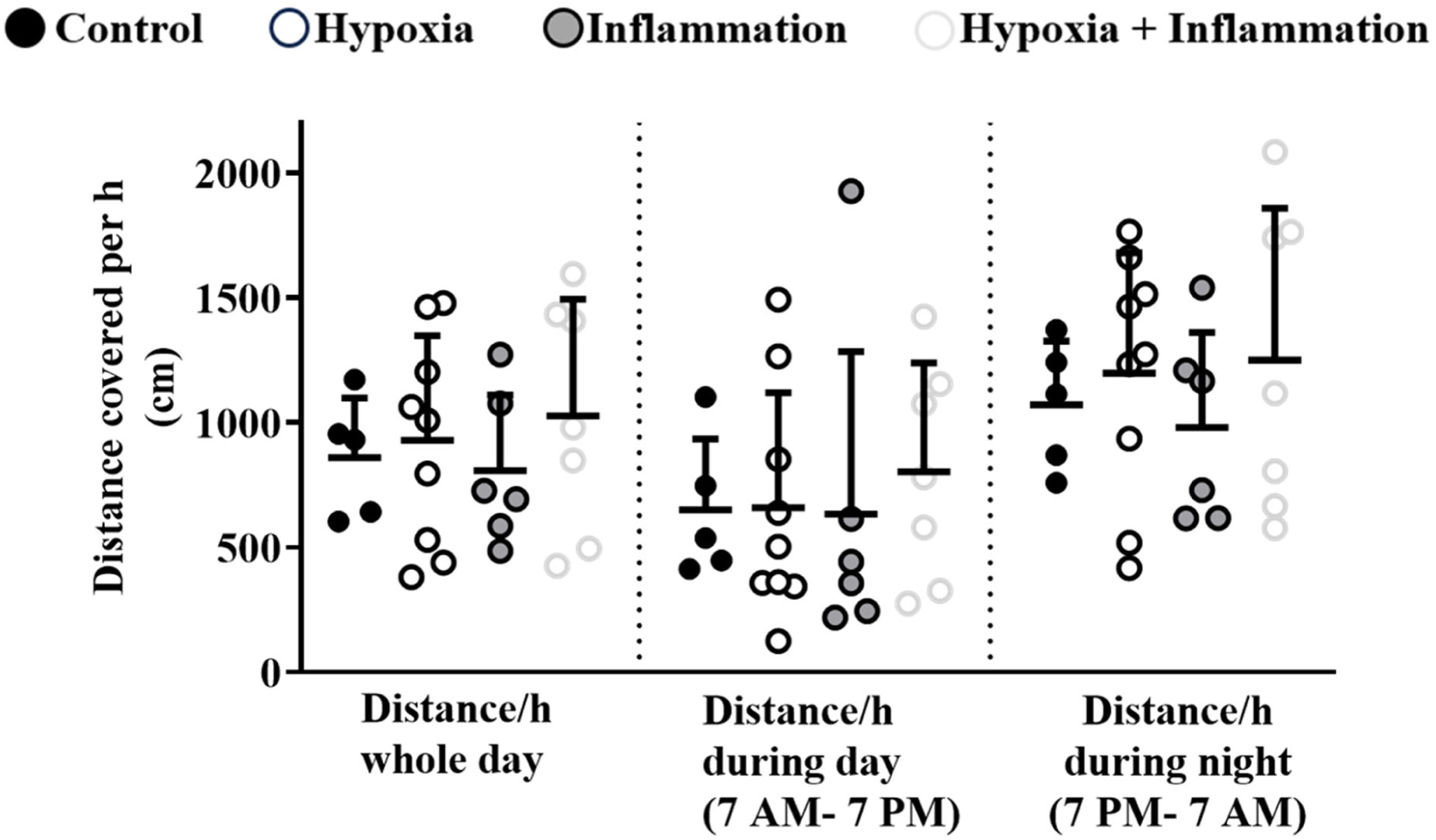
Spontaneous locomotor activity as the distance covered per hour during hypoxia and/or inflammation exposure determined by 24-hour videotracking recording. Circles indicate individual data points (n = 5-9). Horizontal lines represent means and error bars indicate SD. Data were analyzed using 2-way ANOVA.

### Food intake, body and muscle weight

Total food consumption over the study period was lower in animals exposed to hypoxia (NC + I: 672 ± 72.2 g vs H + HI: 472 ± 63.5 g; p < 0.001; η^2^_p_ = 0.76; Supplemental Figure S3A). When adjusted to weekly body weight, we observed that chronic hypoxia led to a transient reduction of food intake during the first two weeks of exposure (1^st^ week: -53%; p < 0.001; η^2^_p_ = 0.59; 2^nd^ week: -24.1%; p < 0.05; η^2^_p_ = 0.48; Supplemental Figure S3B). There were no differences in the starting body weight among the groups (Figure 3A). Nevertheless, rats from the hypoxic groups had significantly lower body weight at the end of the exposure compared to the rats from the normoxic groups (NC + I: 483 ± 51.8 g vs H + HI: 425 + 41.5; p < 0.001; Figure 3A). This was attributed to a reduction in body weight during the first week of the exposure (-10.57%; p < 0.001; η^2^_p_ = 0.72; Figure 3B) as there were no differences during the rest of the study period with similar weekly body weight change. To examine whether chronic hypoxia and pulmonary inflammation led to change in muscle mass, we used the ratio of muscle weight/tibia length as a marker. Pulmonary inflammation led to a significant decline in the mass of both soleus (-11%; p < 0.05; η^2^_p_ = 0.115; Figure 3C) and plantaris (-8.4%; p < 0.01; η^2^_p_ = 0.185; Figure 3D). In contrast, chronic hypoxia decreased the weight of plantaris (-9.11%; p < 0.001; η^2^_p_ = 0.223; Figure 3D) but not that of soleus (Figure 3C).

**Figure 3.**
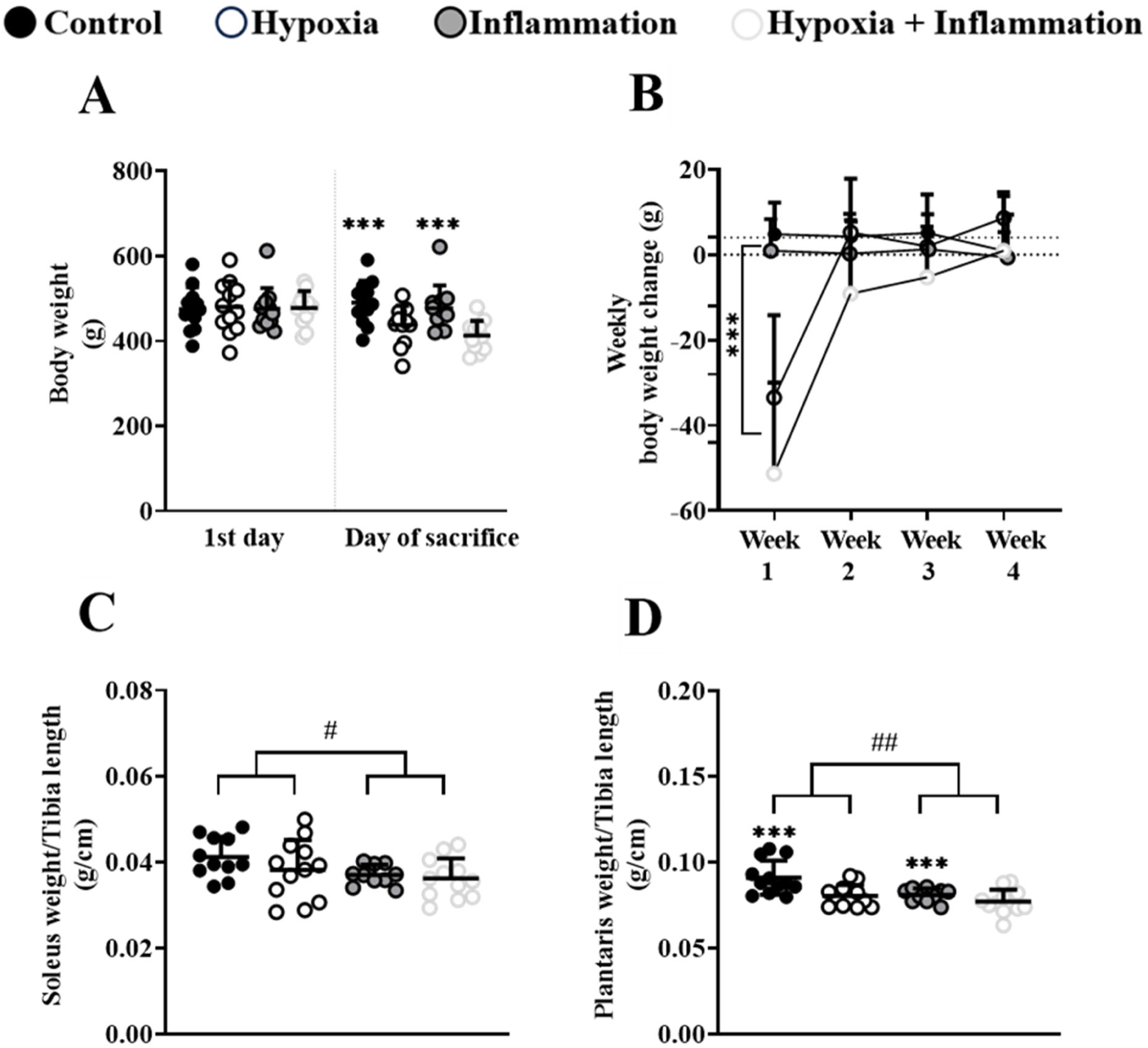
Body and muscle weight under hypoxia and/or pulmonary inflammation conditions. **A**: Starting and final body weight. **B**: Weekly changes in the body weight of the rats throughout the 4-week study period. **C**: Soleus weight at sacrifice normalized to tibia length. **D**: Plantaris weight at sacrifice normalized to tibia length. In **A**, **C** and **D**, circles indicate individual data points (n = 10-12). Horizontal lines represent means and error bars indicate SD. In **B**, circles indicate means and error bars represent SD. Data were analyzed using 2-way ANOVA. ***: p < 0.001, global effect of hypoxia (NC + I vs H + HI). ##: p < 0.01, global effect of inflammation (NC + H vs I + HI). #: p < 0.05, global effect of inflammation (NC + H vs I + HI).

### Fulton index

To validate the physiological efficacy of the hypoxic exposure, we examined the Fulton index as defined above. We observed that hypoxia led to a significant increase in Fulton index (+ 75.23%; p < 0.001; η^2^_p_ = 0.76; Supplemental figure S4A) because of right ventricle hypertrophy (+ 64.7%; p < 0.001; η^2^_p_ = 0.64; Supplemental Figure S4B).

### Mitochondrial oxygen consumption

Measurement of oxygen consumption rate by isolated muscle mitochondria revealed that when GM was used as a substrate, basal (state 2) respiration was significantly lower in the soleus from rats subjected to pulmonary inflammation (-51.7%; p < 0.05; η^2^_p_ = 0.23; Figure 4A) as well as in ADP-stimulated (state 3) respiration (-53.4 %; p < 0.01; η^2^_p_ = 0.33; Figure 4A). Neither pulmonary inflammation nor hypoxia changed CI-driven leak (state 4) respiration (Figure 4A). Mitochondrial respiration in soleus was not altered by inflammation or hypoxia when S (Figure 4B) or GMS (Figure 4C) fueled the electron transport chain. For plantaris, neither pulmonary inflammation nor hypoxia influenced mitochondrial respiration regardless of the substrate used (Figure 4D-F). Finally, respiratory control ratios (State 3/State 4) were unaltered following exposure to inflammation and/or hypoxia (Supplemental Figure S5).

**Figure 4.**
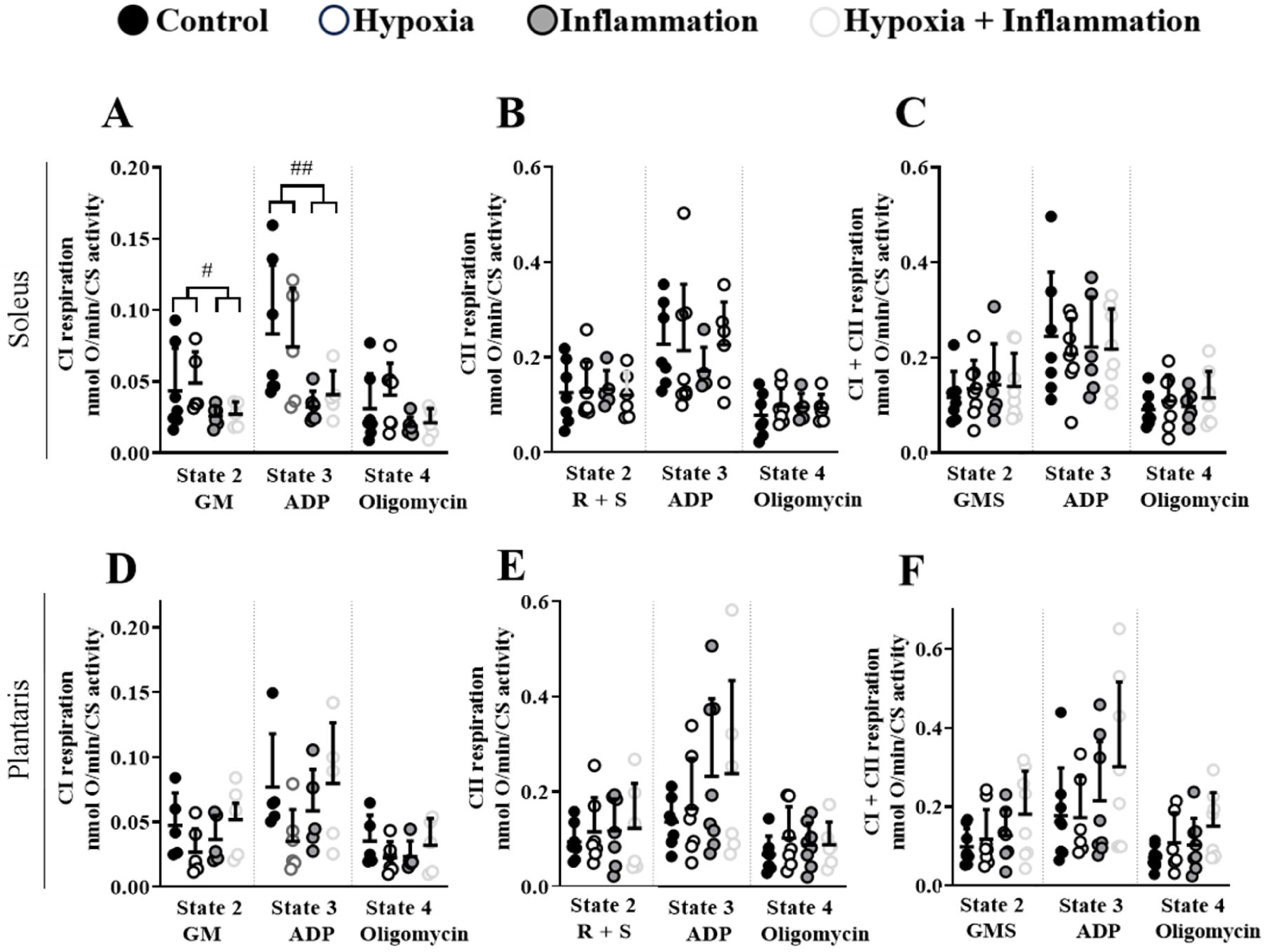
Oxygen consumption in isolated mitochondria following exposure to hypoxia and/or inflammation. Soleus and plantaris mitochondrial respiration measured with Glutamate+Malate (GM) (**A, D**), Rotenone+Succinate (R+S) (**B, E**) and Glutamate+Malate+Succinate (GMS) (**C, F**) substrates, before (basal state 2 condition), after ADP addition (ADP-stimulated state 3 condition) or after addition of oligomycin (leak state 4 condition). All data are normalized to CS activity. Soleus data are presented in panels **A-C** while plantaris data are shown in panels **D-F**. Circles indicate individual data points (n = 5-8). Horizontal lines represent means and error bars indicate SD. Data were analyzed using 2-way ANOVA. **#:** p < 0.05, global effect of inflammation (NC + H vs I + HI); **##:** p < 0.01, global effect of inflammation (NC + H vs I + HI).

### Protein expression

To assess potential changes in mitochondrial protein expression, we examined the protein content of representative subunits of CI and ATP synthase (Complex V, CV) alongside OPA1, a key regulator of the inner mitochondrial membrane fusion. We observed that pulmonary inflammation led to a decrease in the protein content of CV (-38.29%; p < 0.05; η^2^_p_ = 0.192; Figure 5A-B) in soleus. In addition, pulmonary inflammation led to a significant increase in CI/CV ratio (+ 59%; p < 0.01; η^2^_p_ = 0.334; Figure 5A-B). Protein content of CI and CV in plantaris were not influenced by pulmonary inflammation or chronic hypoxia (Figure 5C-D).

**Figure 5.**
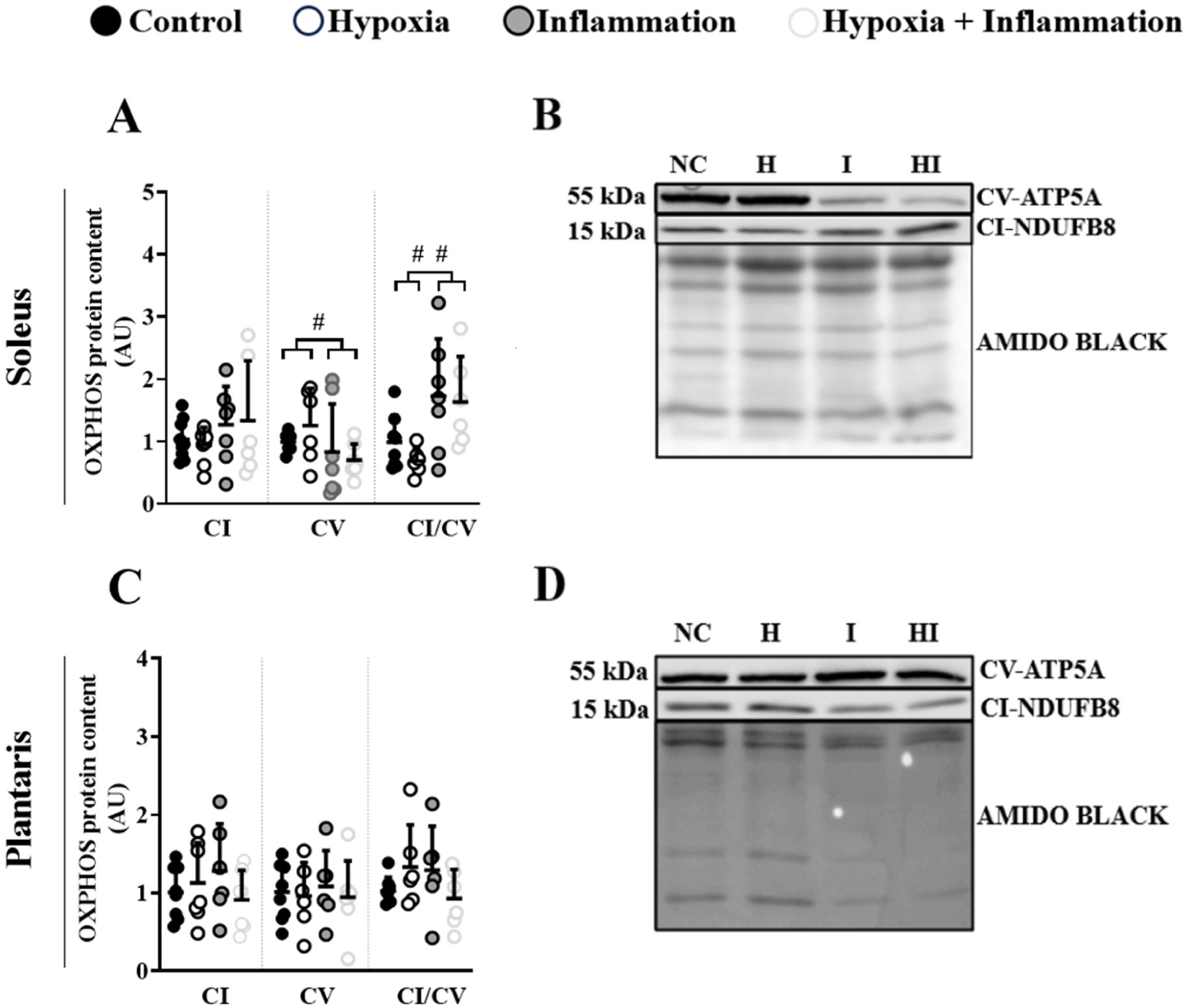
CI and CV protein content and CI-to-CV ratio following exposure to hypoxia and/or inflammation. A &. **C**: Protein content of CI and CV, and CI/CV ratio determined from Western blots on muscle homogenates. **B** & **D**: Representative blots for CI and CV. Amido black staining served as loading control. Soleus data are shown on panels **A** and **B**. Plantaris data are shown on panels **C** and **D**. Circles indicate individual data points (n = 6-9). Horizontal lines represent means and error bars indicate SD . Data were analyzed using 2-way ANOVA. #: p < 0.05, global effect of inflammation (NC + H vs I + HI); ##: p < 0.01, global effect of inflammation (NC + H vs I + HI).

In soleus, pulmonary inflammation resulted in a decrease in the content of total OPA1 (-83%; p < 0.001; η^2^_p_ = 0.519; Figure 6A-B), long form of OPA1 (L-OPA1, -82.5%; p < 0.05; η^2^_p_ = 0.246; Figure 6A-B) and short form of OPA1 (S-OPA1, -92.8%; p < 0.05; η^2^_p_ ^=^ 0.331; Figure 6A-B). However, no differences were observed for the ratio of L - OPA1 to S - OPA1 (L/S, Figure 6A). In contrast with soleus, neither hypoxia nor inflammation changed the protein levels of total OPA1, L-OPA1, S-OPA1 or the L/S ratio in plantaris (Figure 6C-D).

**Figure 6.**
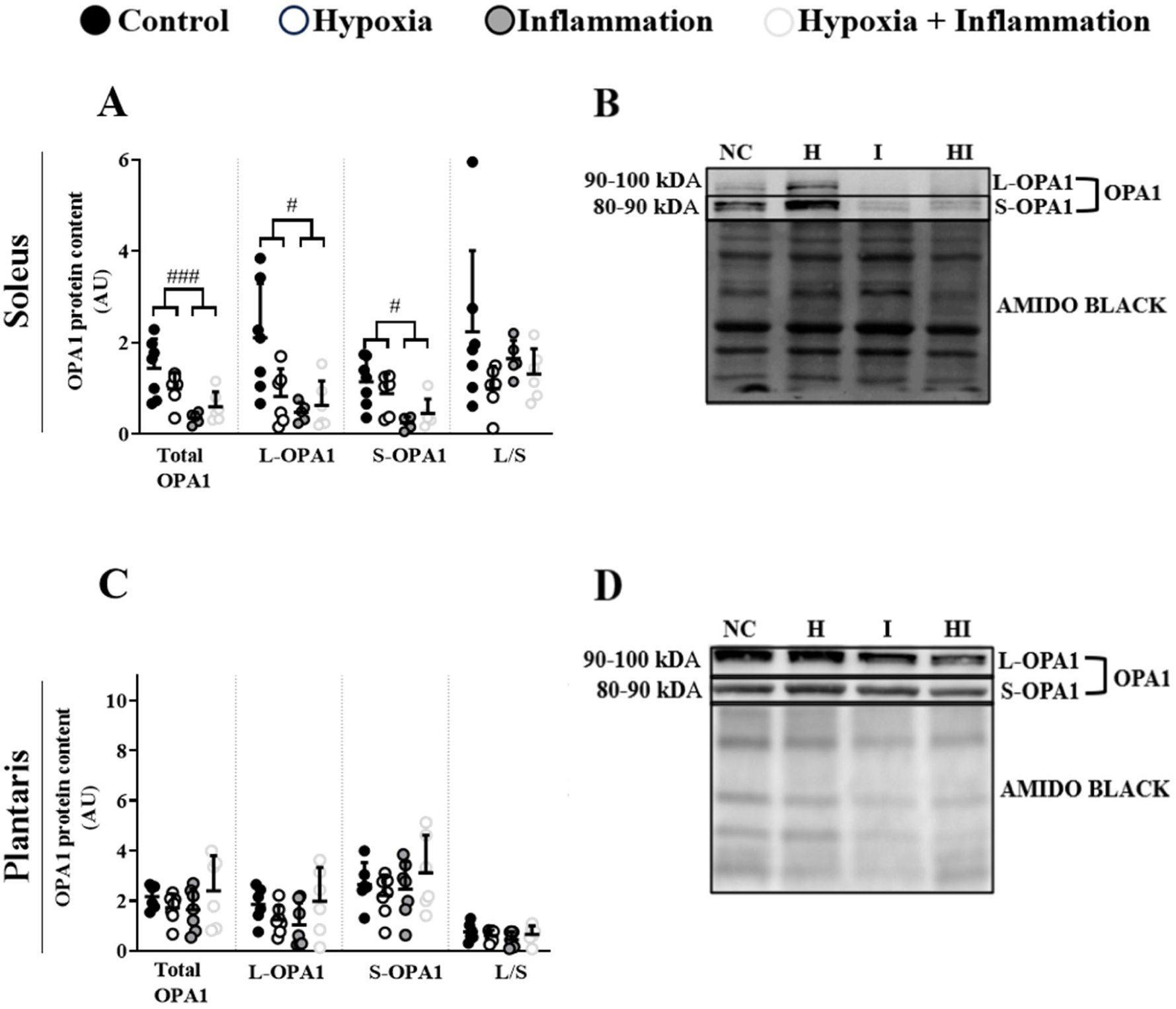
Protein content of OPA1 following exposure to hypoxia and/or inflammation. L-forms and S-forms of OPA1 were identified between 80-100 kD. **A, C:** Protein content of total OPA1, L-OPA1, S-OPA1 and L/S ratio determined by Western blots on muscle homogenates. **B**, **D**: Representative blots for OPA1 in soleus (**B**) and plantaris (**D**). Amido black staining served as loading control. Circles indicate individual data points (n = 5-7). Horizontal lines represent means and error bars indicate SD. Data were analyzed using 2-way ANOVA. #: p < 0.05, global effect of inflammation (NC + H vs I + HI); ###: p < 0.001, global effect of inflammation (NC + H vs I + HI).

### Enzymatic activities

Given the alterations in the CI-supported oxidative phosphorylation of soleus, we further examined whether there were any differences in the enzymatic activity of CI in the same muscle. Pulmonary inflammation led to a significant decrease in CI activity in the two inflammatory groups when compared to normoxia control conditions (I: -97.4%, p < 0.001; HI: -64.3%, p < 0.05; Figure 7A). Chronic hypoxia alone also caused a significant decline in CI activity in soleus muscle (H: -65.9%, p < 0.05; Figure 7A).

To understand whether this decrease in CI activity may have been attributed to a decline in mitochondrial content, we further assessed CS activity in soleus. There were no alterations in CS activity in response to either stressor in soleus muscle (Figure 7B). The same result was also observed in plantaris (Figure 7D).

Due to the decrease in ATP synthase expression in soleus (Figure 5A-B), we further assessed its hydrolytic activity to explore potential functional implications. Nevertheless, we observed that there were no significant effects of inflammation or hypoxia on the isolated hydrolytic activity of ATP synthase in soleus (Figure 7C).

Neither inflammation nor hypoxia alone affected HAD activity in soleus. However, their combination decreased HAD activity when compared with either hypoxia (-61.7%, p < 0.001; Supplemental Figure S6A) or inflammation alone (-62%, p < 0.001; Supplemental Figure S6A). In contrast, there were no significant differences when compared to the control (p = 0.065; Supplemental Figure S6A). In plantaris neither condition altered HAD activity (Supplemental Figure S6A). Finally, LDH activity remained unchanged following hypoxia and/or inflammation in both muscles (Supplemental Figure S6B).

**Figure 7.**
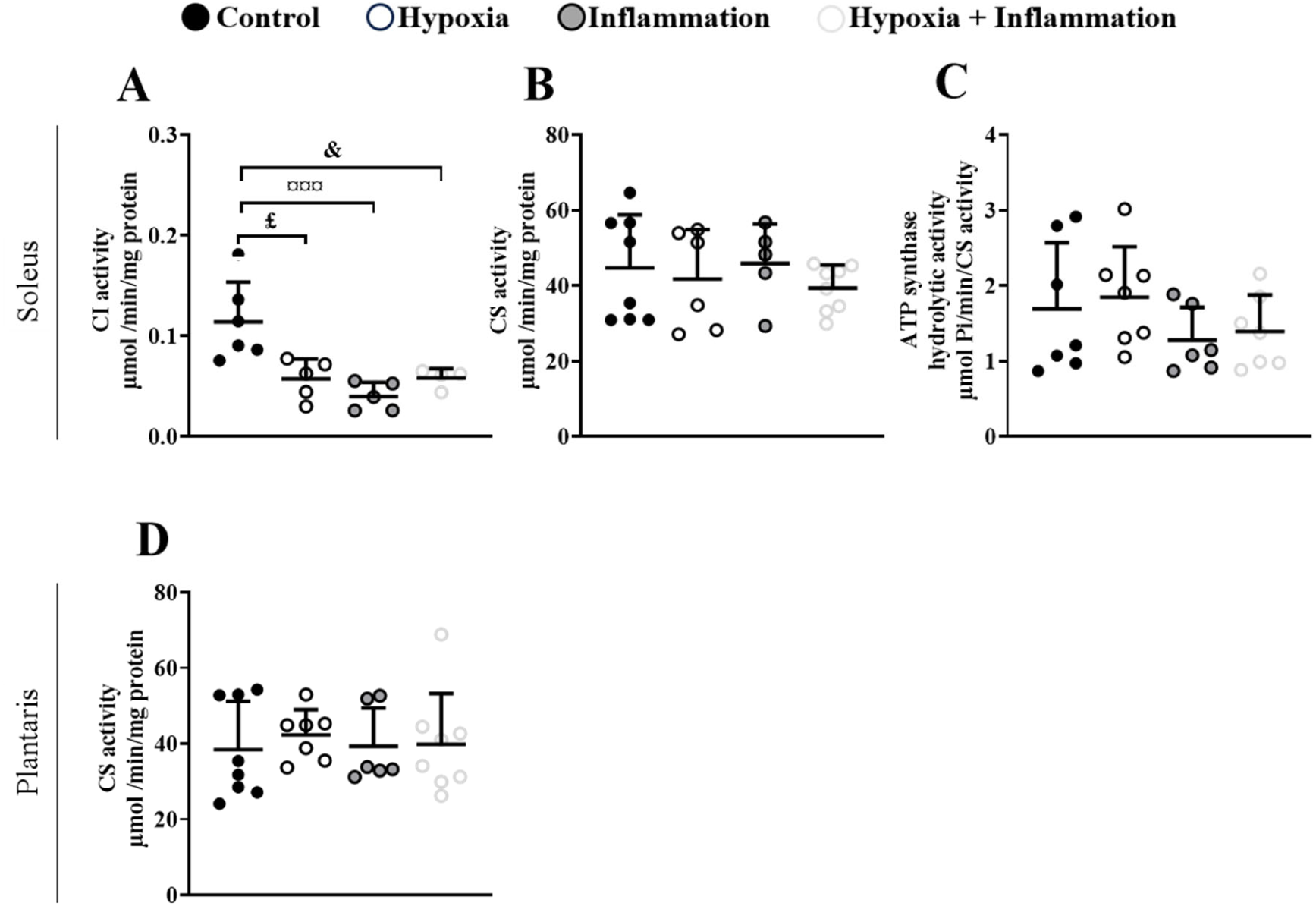
Mitochondrial enzyme activities following exposure to hypoxia and/or inflammation. CI activity was measured in whole soleus homogenates (**A**). CS activity was measured in whole muscle homogenates from soleus (**B**) and plantaris (**D**). ATP synthase hydrolytic activity was measured in isolated mitochondria from soleus (**C**) and normalized to CS activity. Circles indicate individual data points (n = 4-7). Horizontal lines represent means and error bars indicate SD. Data were analyzed using 2-way ANOVA. £: p < 0.05, NC vs H; ¤¤¤: p < 0.001, NC vs I; &: p < 0.05, NC vs HI.

### Regulation of catabolic and anabolic signalling

There was a reduction in pAkt^ser473^/Akt ratio in soleus muscle in response to both pulmonary inflammation (-59.5%, p < 0.01, η^2^_p_ = 0.264; Figure 8A & B) and hypoxia (-61.1 %, p < 0.01, η^2^_p_ = 0.335; Figure 8A & B). In contrast, neither inflammation nor hypoxia affected the pAkt^ser473^/Akt ratio in plantaris following (Figure 8C & D). Neither condition altered REDD1 protein content in soleus or plantaris muscle (Figure 8A-D).

**Figure 8.**
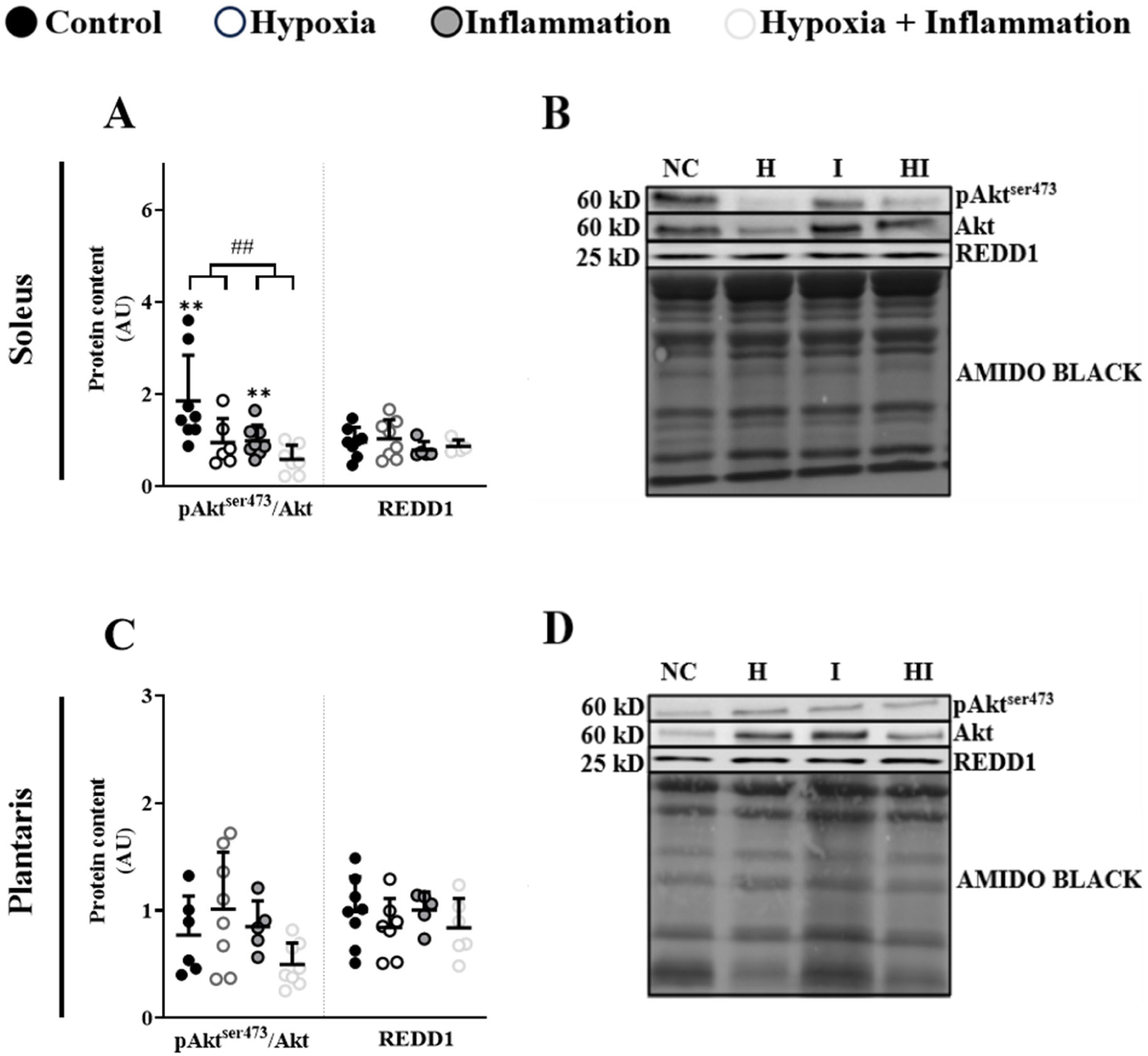
Akt activation and REDD1 protein expression following exposure to hypoxia and/or inflammation. Ratio of pAkt^ser473^ to total Akt and REDD1 protein content determined from Western blots on muscle homogenates in soleus (**A**) and plantaris (**C**) muscles. Representative blots of Akt phosphorylated at serine 473, total Akt and REDD1 in soleus (**B**) and plantaris (**D**). Circles indicate individual data points (n = 5-8). Horizontal lines represent means and error bars indicate SD. Data were analyzed using 2-way ANOVA. ##: p < 0.01, global effect of inflammation (NC + H vs I + HI). **: p < 0.01, global effect of hypoxia (NC + I vs H + HI).

In soleus muscle, inflammation alone (I group) increased the mRNA expression of Atrogin-1 compared to any of the other three groups (+ 56.1% vs NC, p < 0.001; + 76.6% vs H, p < 0.001; + 69.9% vs HI, p < 0.001; Figure 9A). In plantaris, hypoxia alone (H group) increased the mRNA content of Atrogin-1 when compared to the NC group (+93%; p < 0.05; Figure 9C). Nevertheless, there were no effects of inflammation and/or hypoxia on the mRNA content of MuRF-1 neither in soleus nor in plantaris (Figure 9B & D).

**Figure 9.**
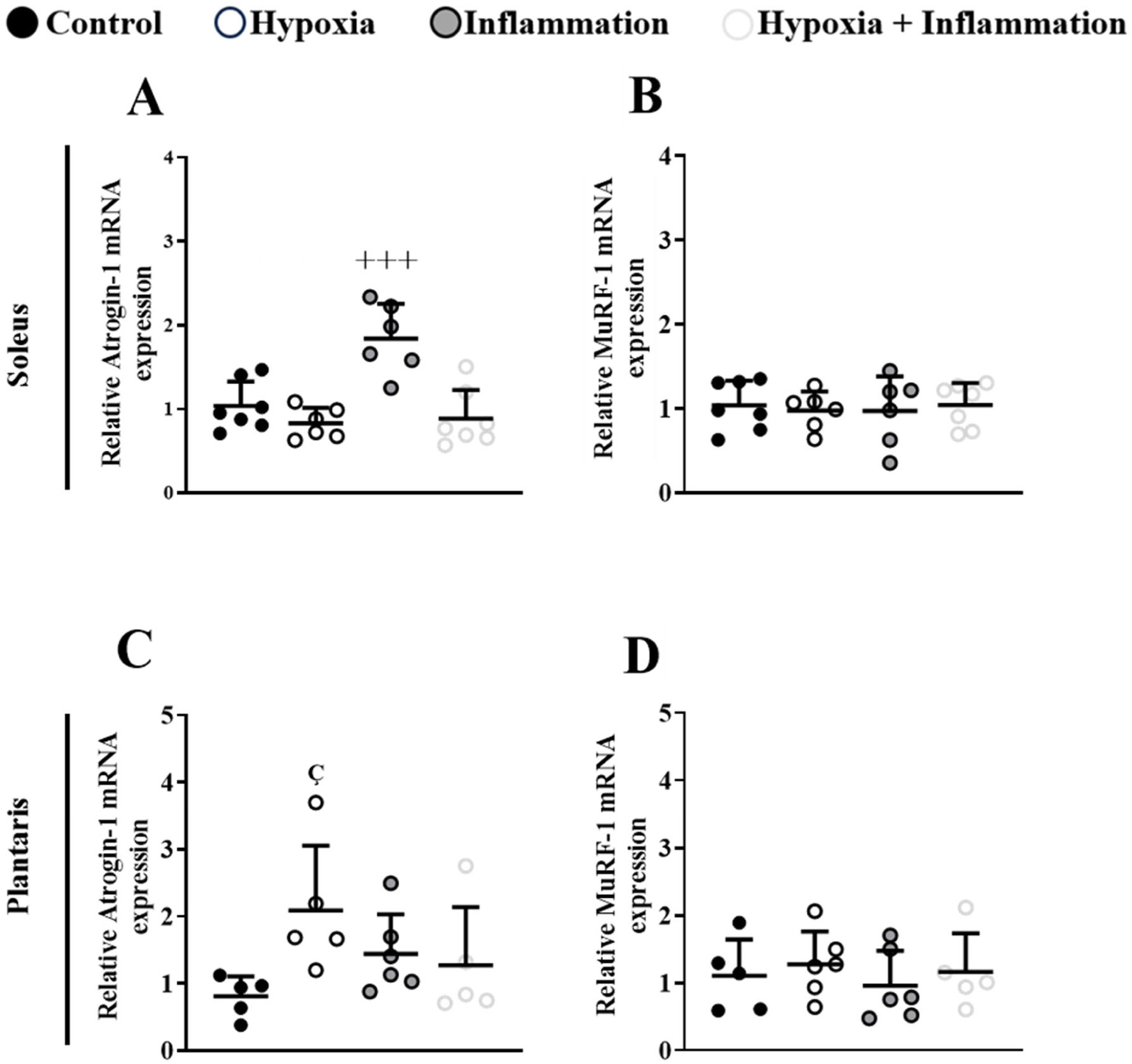
Relative mRNA expression of Atrogin-1 and MuRF-1 gene expression following exposure to hypoxia and/or inflammation. Relative Atrogin-1 and MuRF-1 mRNA expression (normalized to HRPT) in soleus (**A,B**) and plantaris (**C,D**) determined by qPCR. Circles indicate individual data points (n = 5-7). Horizontal lines represent means and error bars indicate SD. Data were analyzed using 2-way ANOVA. **+++**: p < 0.001, I higher than NC, H, HI; ç: p < 0.05, NC vs H.

### Between-cohort analysis

We observed that some of the findings of the present study were in contrast with those from our recently published study (12). Specifically, pulmonary inflammation produced a more pronounced soleus and plantaris phenotype in the present cohort, whereas chronic hypoxia predominantly affected plantaris mitochondrial function in our previous cohort. The intriguing discrepancies between the two studies are summarized in Table 1. Given the divergent responses observed between the two studies, we next compared several physiological and cellular characteristics between the control groups of the two cohorts. Despite being of the same age and strain and obtained from the same supplier, the two cohorts differed considerably in initial body weight, body growth rate, soleus and plantaris mass and plantaris mass-to-body weight ratio (Table 2) In addition, there were significant differences in soleus fiber typology between the two cohorts (Table 2, Supplemental Figure S7). Notably, we observed that in the soleus of the control rats from the current study, there were no hybrid (I/IIa) fibers. In contrast, the hybrid fiber percentage in the soleus of the control group from our previous study was 1.1%. Finally, the differences in body growth between the two rat cohorts were not accompanied by differences in weekly relative food intake (Table 2). We next examined whether physiological traits that differed between the two cohorts were associated with the divergent muscle responses to pulmonary inflammation and hypoxia. We observed that initial body weight was positively associated with soleus mass preservation following exposure to pulmonary inflammation across both cohorts with the corresponding linear regression explaining 38.4% of the variance (Figure 10A). Likewise, a positive association was also observed between initial body weight and relative plantaris mass preservation in inflammatory animals with the corresponding linear regression explaining 25.5% of the variance (Figure 10B).

**Figure 10.**
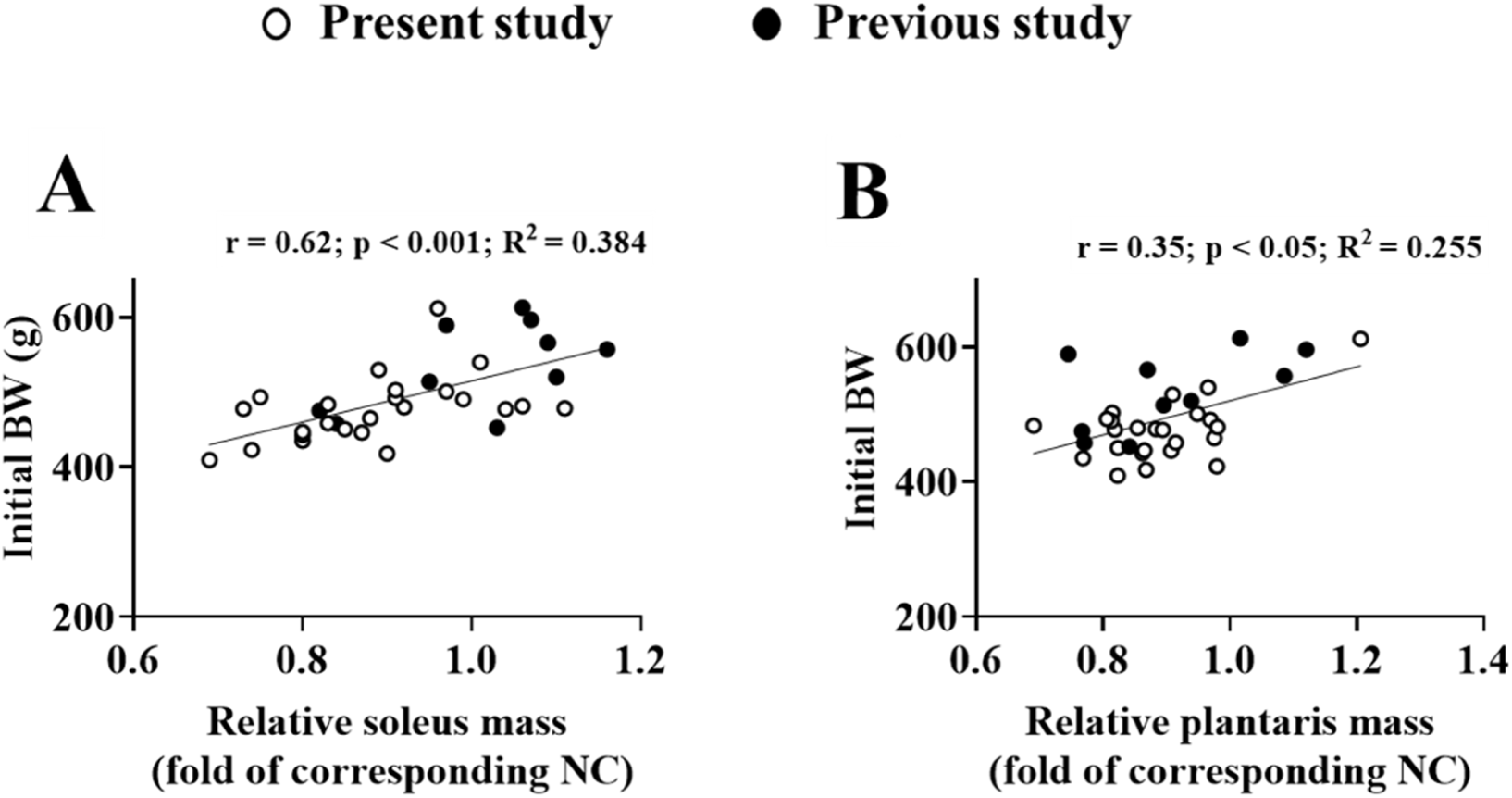
Association between initial body weight and relative muscle mass following exposure to pulmonary inflammation. **A**: Relative soleus mass was calculated by dividing the soleus mass of rats exposed to pulmonary inflammation by the mean soleus mass of the corresponding normoxia control group (n = 34). The association between initial body weight and relative soleus mass was assessed using Pearson’s correlation and linear regression **B:** Relative plantaris mass was calculated using the same approach (n = 34). The association between initial body weight and relative plantaris mass was assessed using Spearman’s correlation and linear regression. White circles represent animals from the present study whereas black circles represent animals from our previous study. The solid line represents the linear regression fit.

**Table 1.** Divergent responses to pulmonary inflammation and hypoxia across the two studies Outcome Present study Previous study.

| Outcome | Present study | Previous study |
| --- | --- | --- |
| <u>Pulmonary inflammation in soleus:</u> |  |  |
| CI-mediated state 3 respiration | Decrease | No change |
| CI activity | Decrease | No change |
| Muscle mass | Decrease | No change |
| <u>Pulmonary inflammation in plantaris:</u> |  |  |
| Muscle mass | Decrease | No change |
| <u>Hypoxia in plantaris:</u> |  |  |
| CI-mediated state 3 respiration | No change | Decrease |

**Table 2.** Differences in physiological and fiber typology between the two rat cohorts.

| Physiological characteristics |  | Present study<br>(n = 6-12) |  | Previous study<br>(n = 6-7) | p value |
| --- | --- | --- | --- | --- | --- |
| <u>Morphological parameters:</u> |  |  |  |  |  |
| Initial body weight | (g) | 477 | ± 49 | 530.8 ± 41 | p < 0.001 |
| Body growth rate | (%) | 3.1 | ± 0.5 | 6.2 ± 1.1 | p < 0.05 |
| Soleus mass | (g) | 0.187 | ± 0.01 | 0.224 ± 0.01 | p < 0.05 |
| Soleus mass/body weight | (mg/g) | 0.38 | ± 0.04 | 0.039 ± 0.02 | NS |
| Plantaris mass | (g) | 0.4 | ± 0.01 | 0.52 ± 0.03 | p < 0.01 |
| Plantaris mass/Body weight | (mg/g) | 0.82 | ± 0.09 | 0.95 ± 0.06 | p < 0.01 |
| <u>Food intake parameters:</u> |  |  |  |  |  |
| Relative food intake week 1 | (g/g/wk) | 0.35 | ± 0.05 | 0.37 ± 0.03 | NS |
| Relative food intake week 2 | (g/g/wk) | 0.37 | ± 0.05 | 0.35 ± 0.02 | NS |
| Relative food intake week 3 | (g/g/wk) | 0.38 | ± 0.08 | 0.35 ± 0.02 | NS |
| Relative food intake week 4 | (g/g/wk) | 0.35 | ± 0.02 | 0.36 ± 0.02 | NS |
| <u>Soleus typology:</u> |  |  |  |  |  |
| Soleus type I fibers | (%) | 99 | ± 1.4 | 93.5 ± 5.1 | p < 0.05 |
| Soleus type IIa fibers | (%) | 1 | ± 1.4 | 5.4 ± 4.4 | p < 0.05 |

For initial body weight, all animals from each cohort were included, as this measurement was obtained before exposure to hypoxia and/or pulmonary inflammation (present study, n = 48; previous study, n = 28). Relative food intake was calculated as food consumption normalized to body weight for each respective week. Muscle mass/body weight ratios are expressed as $\times 10^3$ . Between-cohort differences were assessed using independent-samples t-tests or Mann–Whitney U tests. Values are presented as mean ± SD.

## DISCUSSION

Our findings demonstrate stress- and muscle-specific responses to pulmonary inflammation and hypoxia. Pulmonary inflammation predominantly affected the oxidative soleus inducing atrophy and blunted Akt signaling alongside mitochondrial alterations characterized by impairments in CI function and an increased CI-to-CV protein ratio. These changes were accompanied by a reduction in OPA1 and ATP synthase (Complex V) protein content. In contrast, the glycolytic plantaris did not exhibit mitochondrial alterations following inflammation, despite a decrease in its mass. Chronic hypoxia led to a reduction in CI activity in soleus and induced atrophy only in plantaris, while the combination of inflammation and hypoxia did not result in additive effects for most parameters. These stress- and muscle-specific responses appeared independently of spontaneous locomotor activity that was similar in all experimental groups. Notably, our between-cohort analysis revealed that the physiological phenotype may be associated with differential skeletal muscle responses to chronic pulmonary inflammation and hypoxia.

### Specific effects of pulmonary inflammation

The lack of differences in spontaneous locomotor activity and food intake between inflammatory and non-inflammatory groups indicates that the observed alterations in mitochondrial function and mass of soleus are primarily driven by pulmonary inflammation itself. Furthermore, the absence of changes in CS activity in soleus further suggests that such effects represent impairments in mitochondrial function rather than a reduction in mitochondrial content.

Our results demonstrate atrophy and compromised CI-supported oxidative phosphorylation (OXPHOS) in soleus and align with previous data showing that chronic LPS administration triggers muscle wasting (35) and decline in mitochondrial respiration (36,37). In the current study, the simultaneous reduction in CI activity and CI-driven state 2 and state 3 respiration in soleus points towards a CI dysfunction under various energetic demands. In contrast, unaltered CI-mediated state 4 respiration suggests maintenance of the membrane integrity and mitochondrial potential under non-phosphorylating conditions. Therefore, our data indicate that CI-dependent electron transfer may be limited under increased electron flux.

Inflammation also diminished ATP synthase protein content, leading to altered stoichiometric balance between CI and the phosphorylation system. As the hydrolytic activity of ATP synthase was unaffected, it is likely that the catalytic activity of the enzyme remains functionally intact. This suggests that the reduction in ATP synthase content may not reflect a loss of intrinsic function, but rather a quantitative adjustment to changes in mitochondrial energetics.

Exposure to pulmonary inflammation also elicited a reduction in total OPA1 protein content, affecting both its long (L-OPA1) and short (S-OPA1) forms, without altering their balance. L-OPA1 is primarily involved in maintenance of mitochondrial fusion, cristae integrity and apoptotic resistance while S-OPA1, contributes to cristae remodeling and respiratory efficiency (38). Additionally, a decreased L/S ratio has been associated with membrane depolarization that eventually triggers impaired fusion and unopposed mitochondrial fission (38,39). Although the preserved L/S ratio argues against collapse of the membrane potential and increased mitochondrial fragmentation, the decrease in the abundance of both forms may represent a dysregulation in OPA1 homeostasis. However, as for ATP synthase, the underlying mechanisms of this reduction remain elusive. Furthermore, given the role of OPA1 in cristae stabilization, this alteration may contribute to the observed impairment in CI-dependent respiration (40).

The lack of changes in LDH activity, assessed as the rate of pyruvate reduction to lactate, while it is not directly measuring the glycolytic flux, is consistent with the absence of substantial metabolic shift towards glycolysis to compensate for the blunted mitochondrial function in soleus. This could have contributed to an energy constraint and subsequent downregulation of energy-demanding processes such as protein synthesis. This interpretation is in line with the decline in pAkt^ser473^/Akt in response to pulmonary inflammation implying blunted anabolic signaling through the Akt/mTOR pathway (41). Impaired Akt activation can also promote Forkhead box O dephosphorylation and subsequent activation of the ubiquitin proteasome system (UPS) (42). This aligns with the increase in Atrogin-1 mRNA expression exclusively in the inflammation group, but it indicates that combination of the two stressors leads to UPS-independent proteolysis or unchanged catabolism.

In contrast to soleus, the predominantly glycolytic plantaris exhibited mitochondrial resilience to inflammation. Nevertheless, exposure to inflammation resulted in decline in plantaris mass. Given the absence of mitochondrial alterations in plantaris, the observed atrophy is less likely to be associated with mitochondrial dysfunction. Additionally, our results suggest that plantaris mass wasting occurred independently of alterations in Akt signaling or MuRF-1 and Atrogin-1 mRNA expression. This raises the possibility that other catabolic pathways (43) and/or dysregulation of alternative proteostatic mechanisms (44) may mediate plantaris atrophy under inflammation.

### Specific effects of chronic hypoxia

Chronic hypoxia led to decreased CI activity in soleus. This decline may represent a response to a more reduced state of the ETC characterized by elevated NADH/NAD^+^ ratio due to the lower O_2_ availability (45). Nevertheless, this reduction in isolated CI activity was not accompanied by impairment of CI-driven OXPHOS. This suggests that the remaining CI activity was still sufficient to sustain CI-dependent respiration. In support of this interpretation, earlier data in isolated brain mitochondria have shown that CI activity can decline substantially before major impairments in respiration take place (46). This is consistent with a threshold concept whereby respiration is compromised only after a marked reduction in enzymatic activity. Similarly, a dissociation between CI activity and CI-mediated respiration has been reported following exercise (47).

Chronic hypoxia led to atrophy only in plantaris, whereas soleus mass was preserved despite diminished Akt activation. This is consistent with our recent study (12). The greater vulnerability of glycolytic muscles to hypoxia has also been observed previously (11) and has been attributed to divergent structural adaptations whereby glycolytic muscles exhibit reduction in fiber size to limit oxygen diffusion distance whereas oxidative muscles enhance capillary contacts (11). From a mechanistic standpoint, the atrophy in plantaris may at least partially involve activation of the UPS given that we observed an increase in Atrogin-1 mRNA expression in the hypoxia group.

### Combined effect of pulmonary inflammation and chronic hypoxia

Parallel exposure to hypoxia and inflammation revealed predominantly non-additive interactions between these two stressors. The only parameter for which the combination produced an additional effect was the decrease in soleus HAD activity that may suggest changes in fatty acid oxidation. In contrast, variables such as soleus Akt activation and CI activity that were independently affected by both stressors, were not further altered under combined conditions. For parameters affected selectively by one stressor, the combination of the two conditions generally led to a stress-specific prevalence rather than synergistic effects. For instance, pulmonary inflammation prevailed over hypoxia on soleus CI-driven state 3 respiration while it suppressed the hypoxia-induced decrease in Atrogin-1 mRNA expression in plantaris. Conversely, hypoxia inhibited the inflammation-mediated increase of Atrogin-1 mRNA expression in soleus. Thus, taken together, in most cases hypoxia and inflammation appear to have opposite effects with the resulting phenotype determined by the predominant stressor.

### Comparative analysis

We observed divergent responses to inflammation and hypoxia between our current and previous cohorts (12). Although the two cohorts consisted of age- and strain-matched rats that were obtained from the same supplier, they differed in several physiological characteristics, including initial body weight, body growth rate, soleus and plantaris mass, plantaris mass relative to body weight and soleus fiber typology. Notably, the difference in body growth rate was not accompanied by differences in relative food intake, indicating that food consumption is not likely to contribute to the distinct physiological traits between the two cohorts. Additionally, the differences in soleus fibre typology between the two cohorts further indicates that this phenotypic divergence extended beyond body and muscle mass to the cellular composition of skeletal muscle. These findings imply that animals from apparently identical experimental cohorts can still exhibit substantial differences in physiological characteristics that may partially influence their subsequent responses to a given stressor.

This possibility is supported by the positive correlation between initial body weight and preservation of both soleus and plantaris mass following exposure to pulmonary inflammation. Importantly, the cohort differences varied across muscles and stressors. Specifically, pulmonary inflammation resulted in a more pronounced soleus phenotype in the current study, whereas chronic hypoxia produced a more pronounced plantaris phenotype in our previous study. Therefore, initial body weight cannot be regarded as a general determinant of skeletal muscle response to inflammation or hypoxia as its association with the muscle phenotype was not consistent across outcome measures, muscles and stressors. This would be even more critical when muscle data are expressed relatively to body weight. Rather, these findings imply that physiological traits may affect susceptibility in a stressor- and muscle-specific manner. In addition, the absence of a consistent association between initial body weight and CI-mediated state 3 respiration suggests that the divergent mitochondrial responses across the two cohorts may have been influenced by molecular, cellular or metabolic determinants that were not assessed in our studies.

The phenomenon of inter-individual physiological heterogeneity in skeletal muscle responses has also been observed in chronic diseases. In COPD, skeletal-muscle dysfunction is reported in approximately 30–40% of patients while the magnitude of the response to exercise rehabilitation can vary considerably among these individuals (48,49) . Although such heterogeneity may reflect differences in disease stage, physical activity levels, nutritional status and other co-existing comorbidities (50), our findings raise the possibility that broader physiological differences may also affect the skeletal muscle vulnerability to disease-related stress. Interestingly, the correlation between soleus mass alterations and initial body weight we observed may be in line with the so-called “obesity paradox,” which describes the association between higher BMI and a more favorable prognosis in COPD patients experiencing acute exacerbation, a condition characterized by a heightened inflammatory state (51). Similarly, several epidemiological studies have reported that body mass is associated with mortality risk (52,53). In this context our analysis provides experimental evidence that an individual’s physiological state may be associated with the extent of skeletal muscle alterations in chronic disease. This consideration is also relevant to animal studies, where replication of strain and age may not necessarily guarantee an identical baseline phenotype and subsequently a homogeneous response to a common pathological stimulus.

### Strengths and limitations

The major strength of this study lies within the experimental design that enabled us to disentangle not only the individual effects of the two stressors but also their interaction. From a translational standpoint, this is particularly relevant considering that both stressors are often present in pathophysiological settings such as respiratory diseases known to affect muscle mass and function. Another strength is the measurement of spontaneous locomotor activity that strengthened our interpretations that the mitochondrial phenotype was not confounded by changes in activity levels. In addition, the assessment of L-OPA1 and S-OPA1 provided mechanistic insight into mitochondrial inner membrane fusion beyond total OPA1 content alone. Furthermore, the effectiveness of the hypoxic exposure was validated through the expected increase in Fulton index.

Besides these strengths, our study has limitations. First, only male rats were included to enable a direct comparison with our previous cohort. However, it remains unknown whether the same skeletal muscles responses would have been observed in female rats. In addition, mitochondrial dynamics were determined indirectly through investigation of the OPA1 protein content and form distribution, without a direct assessment of mitochondrial morphology. Another limitation is the lack of pair-fed control for each hypoxic group, which makes it challenging to define the precise extent to which hypophagia contributed to the observed phenotype. However, the muscle-specific pattern of responses indicates that hypophagia is unlikely to account for all the observed alterations.

## Conclusion

Using an original rodent model, we demonstrate that chronic hypoxia and pulmonary inflammation induce muscle-specific alterations in skeletal muscle mass and mitochondrial function that are independent of locomotor activity levels. Pulmonary inflammation drives atrophy in both muscles but triggers mitochondrial alterations, highlighted by CI dysfunction, primarily in oxidative soleus. In contrast, hypoxia preferentially induces atrophy in the predominantly glycolytic plantaris. Their combination does not exacerbate the specific effects of each independent stressor, but it rather appears to create an antagonistic environment where one factor may obscure the effects of the other. Notably, comparison with our previous cohort revealed that animals from identical experimental models differ considerably with regards to several physiological traits and responses to the same stressors. Consequently, our findings highlight the need to account for animal and muscle phenotypes, as well as inter-individual physiological heterogeneity, when interpreting skeletal muscle responses in experimental models of chronic disease.

## DATA AVAILABILITY

Data will be made available upon reasonable request.

## ACKNOWLEDGMENTS

A.G. was funded a 3-year PhD fellowship from Université Grenoble Alpes (IRGA-2022 EPIMUS) and experiments were funded by Fondation Agir Pour les Maladies Chroniques (RAK20007CCA). We acknowledge the LBFA Imaging facility, member of Grenoble-IsDV IBISA labelled perimeter (Grenoble Imagerie Sciences du Vivant) and member of the national infrastructure France-BioImaging supported by the French National Research Agency (ANR-10-INBS-04-01).

## AUTHOR CONTRIBUTIONS

A.G, C.Ch and H.D conceived and designed research; A.G, O.L. N.B, and C.T performed experiments; C.Co performed microscopy imaging; A.G analyzed data; A.G, C.Ch and H.D interpreted results of experiments; A.G prepared figures and drafted manuscript; A.G, C.Ch and H.D edited and revised manuscript; A.G, O.L, N.B, C.Co, C.T, C. Ch and H.D approved final version of manuscript.

