## Supplemental_figures for "Distinct patterns of skeletal muscle wasting and mitochondrial changes following hypoxia and inflammation: associations with physiological phenotype"

### Supplementary material

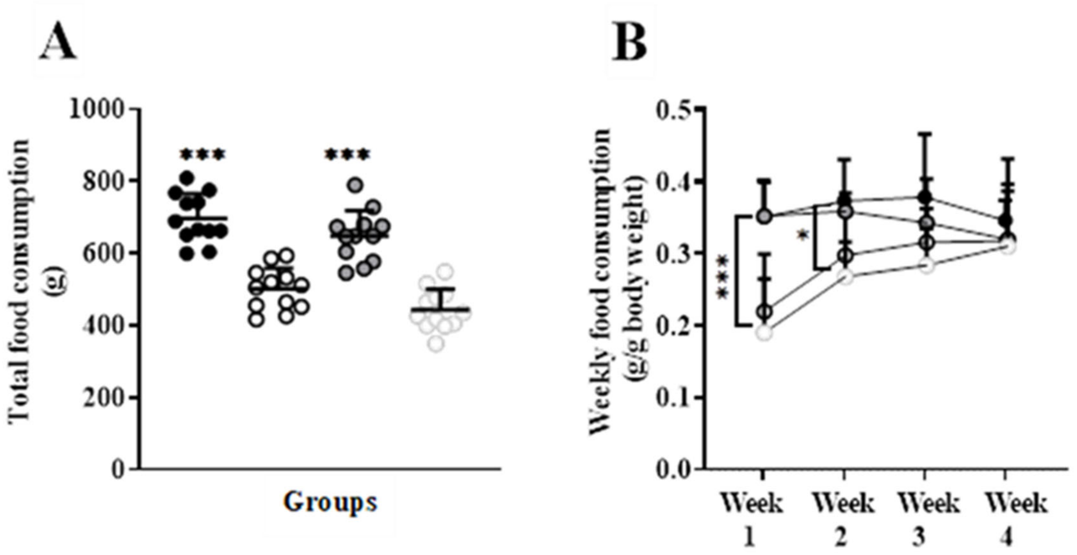


**S3. Food consumption under hypoxia and pulmonary inflammation conditions. A**: Total food consumption over the 4-week period. B: Weekly changes in relative food intake throughout the 4-week exposure. Relative food intake was calculated as food consumption normalized to body weight for each respective week In **A**, circles indicate individual data points (n = 12). Horizontal lines represent means and error bars indicate SD. In **B**, circles represent means and error bars represent SD (n = 12). Data were analyzed using 2-way ANOVA ***: $p < 0.001$, global effect of hypoxia (NC + I vs H + HI). *: $p < 0.05$, global effect of hypoxia (NC + I vs H + HI).

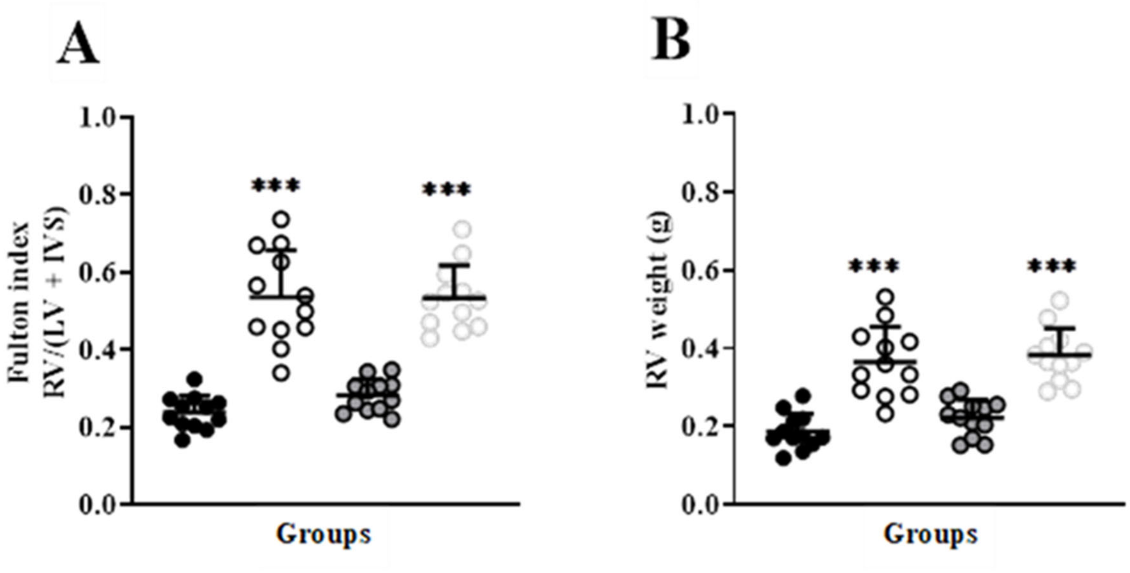


**S4. Fulton index and RV weight following exposure to hypoxia and/or inflammation. A:** Fulton index calculated as the ratio of right ventricle (RV) weight to the combined weight of left ventricle (LV) and intraventricular septum (IVS). **B:** Weight of the RV. Circles indicate individual data points (n = 12). Horizontal lines represent means and error bars indicate SD. Data were analyzed using 2-way ANOVA *****: $p < 0.001$, global effect of hypoxia (NC + I vs H + HI).

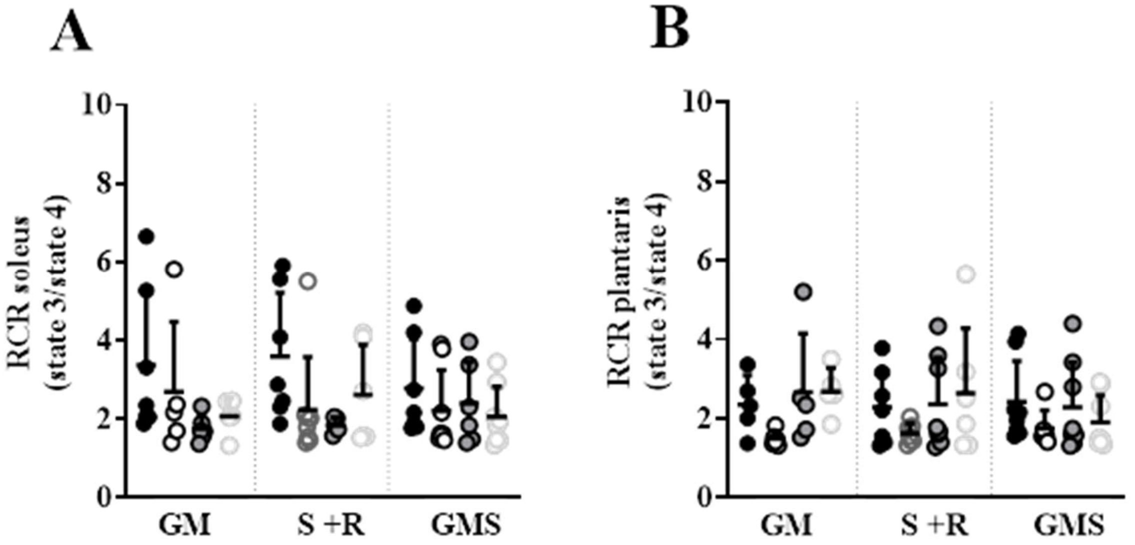


**S5. Respiratory control ratio (RCR) in isolated mitochondria from soleus and plantaris following exposure to hypoxia and/or inflammation.** Soleus (A) and plantaris (B) RCR (state 3/state 4) with CI (GM), CII (S+R) and CI + CII (GMS) substrates. Circles indicate individual data points (n = 5-8). Horizontal lines represent means and error bars indicate SD**.** Data were analyzed using 2-way ANOVA.

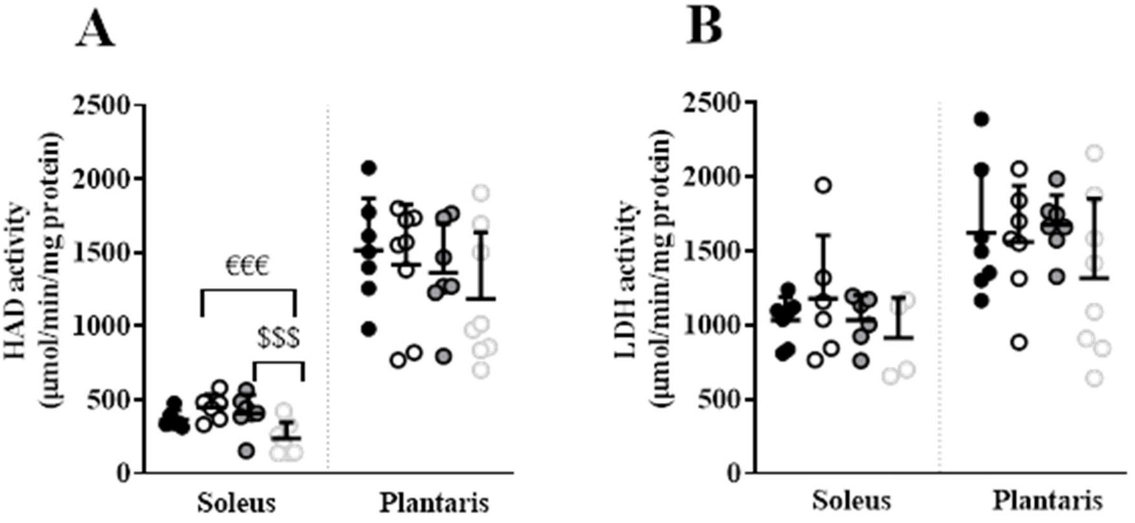


**S6. HAD (A) and LDH (B) activity in whole soleus and plantaris muscle homogenates following exposure to hypoxia and/or inflammation**. Circles indicate individual data points (n = 4-8). Horizontal lines represent means and error bars indicate SD. Data were analyzed using 2-way ANOVA €€€: $p < 0.001$, H vs HI; $$$: $p < 0.001$, I vs HI.

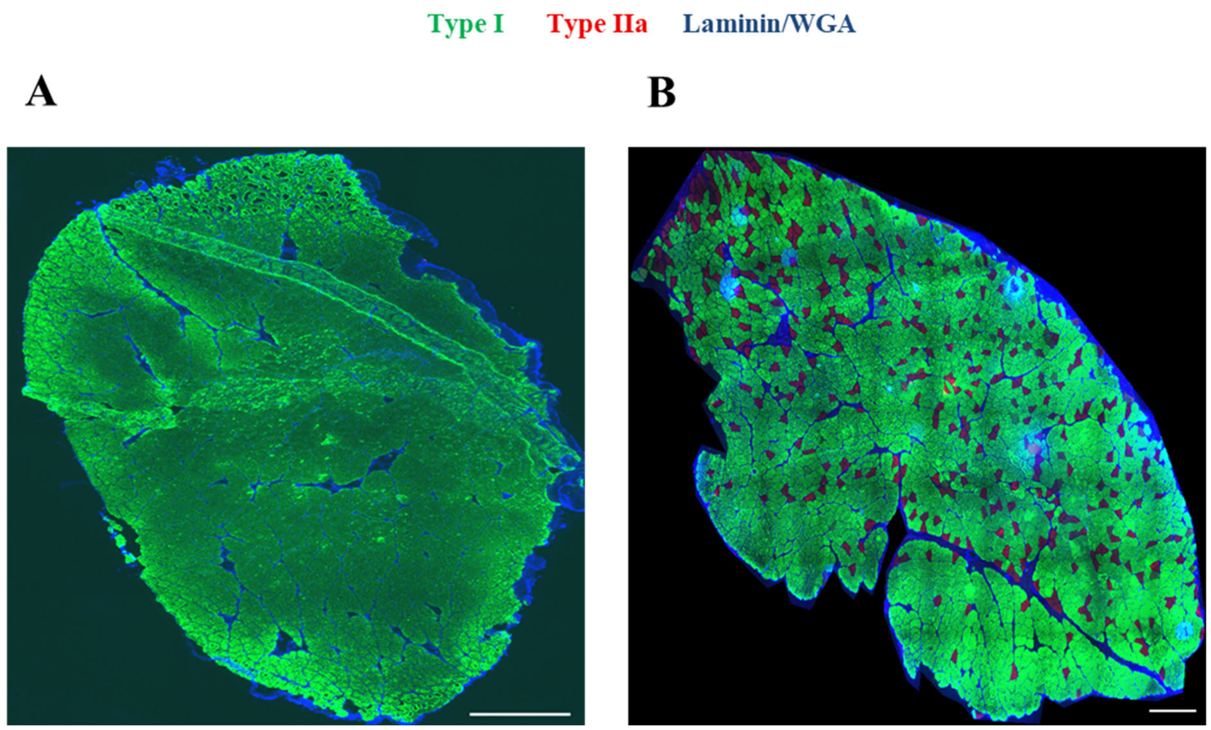


**S7. Soleus fiber typology in controls from the present and previous studies.** Representative whole-soleus cross-sections from the control animals in the present (**A**) and previous (**B**) studies. Tissue sections were processed via multi-channel immunofluorescence staining to identify cell boundaries (laminin in A and wheat germ agglutinin,WGA in B; blue), type I fibers (MHC-I; green), and type IIa fibers (MHC-IIa; red). Blue fluorescent circular aggregates in panel B represent non-specific WGA precipitation artifacts and did not interfere with the identification of myofiber boundaries or fiber-type classification. White scale bars = 0.5 mm.
