## Supplemental_tables for "Distinct patterns of skeletal muscle wasting and mitochondrial changes following hypoxia and inflammation: associations with physiological phenotype"

### Supplementary material

**S1. Primary antibodies used for western blotting.**

| Target | Manufacturer | Product Nr | Dilution |
|---|---|---|---|
| **OXPHOS** | Abcam | ab110413 | 1:1000 |
| **OPA1** | Cell Signaling Technology™ | #80471 | 1:1000 |
| **REDD1** | Santa Cruz Biotechnology | (sc-46034) | 1:1000 |
| **Akt** | Cell Signaling Technology™ | #4691 | 1:1000 |
| **pAkt$^{ser473}$** | Cell Signaling Technology™ | #9271 | 1:2000 |

***Abbreviations:*** OXPHOS, oxidative phosphorylation; OPA1, optic atrophy 1; REDD1, Regulated in DNA Damage and Development 1; Akt, protein kinase B: ser, serine

**S2. Sequences and characteristics of the primers used in this study.**

| Primers | Forward sequence | Forward primers concentration (µM) | Reverse sequence | Reverse primers concentration (µM) | Annealing temperature (°C) |
|---|---|---|---|---|---|
| HRPT | 5'-ggtccattcctatgactgtagatttt- 3' | 0.66 | 5'-caatcaagacgttctttccagt t-3' | 0.66 | 60 |
| MuRF-1 | 5'-agcattgtagaagcttccaagg-3' | 0.66 | 5'-gtaaactcctcctcctcatctgt c-3 | 0.66 | 58 |
| Atrogin-1 | 5'-gcttgtgcgatgttacccaagaa-3' | 0.66 | 5'-tgaaagtgagacggagcagc tct-3' | 0.66 | 58 |

***Abbreviations:*** HRPT, Hypoxanthine-guanine phosphoribosyltransferase; MuRF-1, Muscle Ring Finger 1
